# Wnt signaling and tissue tension reconstitute planar cell polarity *in vitro*

**DOI:** 10.64898/2026.05.01.722304

**Authors:** Soichi Matsuo, Minako Suzuki, Sayuki Hirano, Tetsuhisa Otani, Yusuke Mii

**Affiliations:** Institute for Life and Medical Sciences, Kyoto University, 53 Shogoin Kawahara-cho, Sakyo-ku, Kyoto 606-8507, Japan; Graduate School of Biostudies, Kyoto University, Yoshida-Konoecho, Sakyo-ku, Kyoto, 606-8501, Kyoto, Japan; Department of Biological Sciences, Graduate School of Science, Tokyo Metropolitan University, Hachioji, Tokyo 192-0397, Japan; Department of Chemical Science and Engineering, Graduate School of Engineering, Kyoto University, Kyoto 606-8507, Japan

## Abstract

Planar cell polarity (PCP) aligns cells within the plane of a tissue through asymmetric localization of core PCP proteins. *In vivo*, PCP arises from integration of biochemical signaling and mechanical inputs, making it challendging to reconstruct *in vitro*. Here we reconstitute PCP in cultured MDCK epithelial cells using Wnt signaling, collective migration, and Prickle3 overexpression. Loss of Vangl1 disrupts reconstituted PCP, directional collective migration, and propagation of ERK traveling waves. Orthogonal manipulation of signaling and mechanics show that tissue tension contributes to establishing a polarity axis, but not its direction, whereas local Wnt input specifies polarity direction. Together, these inputs generate tissue-wide vectorial polarity. This system provides a tractable framework for dissecting how signaling and mechanics are integrated to organize tissues.

**Teaser:** Wnt signaling and tissue tension reconstitute planar cell polarity in cultured epithelial cells through core-component interdependence and ERK dynamics.

## Introduction

Synthetic biology seeks to understand biological phenomena by reconstructing them from defined components, including engineered proteins and genetically modified cells. This strategy is now being extended from cell-autonomous processes to multicellular systems, with the goal of understanding how embryonic cells coordinate their behavior and how cell populations organize into higher-order tissue structures. A central feature of embryonic development is positional information, which is often provided by secreted morphogens such as Wnt and BMP. Morphogens are classically thought to form concentration gradients that cells interpret in a dose-dependent manner to infer their position. Supporting this concept, arbitrary diffusible proteins such as GFP and mCherry can be converted into synthetic morphogens by combining localized secretion, extracellular trapping, and synthetic receptors, enabling positional encoding and spatial patterning in engineered mammalian cells (*1*) and in the *Drosophila* wing disc (*2*). In parallel, synthetic cell-cell communication systems have made cell adhesion programmable. For instance, synNotch-induced differential cadherin expression is sufficient to drive multicellular self-organization, symmetry breaking of tissue-like structures, and regenerative rearrangements *in vitro* (*3*). Yet most current synthetic systems reconstitute cell-fate patterning or tissue organization, whereas developmental control systems that integrate signaling, mechanics, and intercellular feedback remain far less tractable *in vitro*.

Planar cell polarity (PCP) is one such system. PCP refers to the coordinated orientation of cells within the tissue plane orthogonal to the apicobasal axis and is essential for a wide range of morphogenetic processes and tissue functions across metazoans (*4–10*). In vertebrates and *Drosophila*, PCP depends on a conserved set of core components, including the transmembrane proteins Frizzled(*11*), Van Gogh-like (Vangl) (Van Gogh or Strabismus in *Drosophila*) (*12, 13*), and Celsr (Flamingo in *Drosophila*) (*14*), and the cytoplasmic proteins Dishevelled (*15, 16*) and Prickle (Pk) (*17*). These proteins adopt characteristic polarized distributions in individual cells: Celsr is typically enriched bipolarly at opposite cell boundaries (*14*), whereas Pk, Frizzled, and Dishevelled are localized unipolarly (*18, 19*), respectively, corresponding to axial and vectorial polarity. Through intercellular complexes, their localization is mutually regulated across cell boundaries, thereby generating robust tissue-scale polarization and morphogenesis (*20–22*). This tissue-scale polarization is linked to a particular body axis, such as the anterior-posterior axis in *Xenopus* neural plate and the proximal-distal axis in Drosophila wing, in all *in vivo* PCP systems.

In addition to the interdependence of core PCP components, vertebrate PCP is controlled by multiple converging inputs, including Wnt signaling, additional context-dependent inputs such as FGF, and tissue mechanics (*23–26*). Noncanonical (β-catenin independent) Wnt signaling has long been implicated in PCP-related morphogenesis. Wnt11 regulates gastrulation and convergent extension movements through a β-catenin-independent pathway during early embryogenesis in *Xenopus* and zebrafish (*27, 28*). Our recent work further suggests that endogenous Wnt11 in the *Xenopus* neural plate does not form an obvious long-range gradient along the axis of planar polarity, but instead accumulates in a planar-polarized manner at cell boundaries together with core PCP components, consistent with formation of intercellular complexes across adjacent cells (*26*). In the mouse limb bud, Wnt5a regulates PCP by promoting Vangl phosphorylation and by enabling FGF-dependent orientation of polarity (*23, 24*). At the molecular level, these inputs converge on the phosphorylation state of Vang/Vangl proteins, which governs their interaction with Prickle and other core factors, thereby influencing assembly and stability of polarized PCP complexes (*29–31*). Consistent with this idea, FGFR1 can modulate PCP in *Xenopus* neuroectoderm via Vangl2 tyrosine phosphorylation (*25*).

Mechanical cues add a further layer of regulation. Tissue strain can orient the planar axis in *Xenopus*, and tissue tension helps maintain and align PCP in the neuroectoderm. Cell flow, tissue stress and shear align PCP in the *Drosophila* wing (*32–35*). Together, these studies indicate that PCP emerges from integration of biochemical and mechanical inputs rather than from any single cue. These observations suggest that PCP reconstitution *in vitro* will require controlled spatial Wnt input together with tissue-level mechanical context. PCP is therefore an attractive target for synthetic reconstruction. However, most current knowledge of PCP comes from *in vivo* systems, and reported *in vitro* PCP models rely on differentiated multiciliated cells that intrinsically acquire planar polarity (*36, 37*). Whether PCP can be reconstituted de novo in a minimal epithelial context therefore remains a fundamental open question.

## Results

### Reconstitution of Wnt11-dependent planar cell polarity *in vitro*

Because adjacent expression of Wnt11 and Pk3 together with Vangl2 is sufficient to induce ectopic planar cell polarity (PCP) in *Xenopus* ectodermal epithelia (*38*), we first asked whether a similar configuration could be recapitulated in MDCK II cells, a widely studied epithelial model. To visualize Pk3, we used mRuby2-Pk3 (RP3) (*26, 34*). In contrast to *Xenopus* (*38*), co-expression of Vangl2 was not required for membrane localization of RP3 (fig. S1A). However, adjacent seeding of Wnt11-overexpressing cells and RP3-overexpressing cells using a two-well chamber failed to induce polarized RP3 localization (fig. S1A).

In *Xenopus* embryos, Pk3 polarization requires not only Wnt11 signaling, but also tissue tension (*34*). Moreover, our recent work suggests that adjacent positioning of high- and low-Wnt regions is sufficient to coordinate PCP over long distances. Therefore, we sought to reconstruct both tissue tension and a spatial contrast of Wnt signaling *in vitro*. In MDCK II cells, removal of a physical confinement after cells reach confluence induces collective migration, in which leader cells at the edge crawl outward and generate tension in the epithelial sheet (*39, 40*). We reasoned that imposing high- and low-Wnt regions during this collective movement would provide the minimal conditions for PCP polarization.

Despite extensive studies on morphogen gradients, methods to artificially control the spatial distribution of secreted proteins remain limited. From our studies in *Xenopus*, we found that binding of secreted proteins to the cell surface is essential for their spatial distribution (*41*). Accordingly, we examined how secreted mVenus distributes in a collectively migrating MDCK epithelial sheet expressing morphotrap, a membrane-tethered anti-GFP nanobody. mVenus accumulated strongly at the leading edge, whereas it exhibited weaker and relatively uniform distribution in the interior (fig. S1B). Cells that lack morphotrap expression showed no detectable mVenus accumulation (fig. S1B), indicating that mVenus distribution depends on cell-surface binding, as in *Xenopus* (*41*). These results indicate that the leading edge efficiently captures secreted proteins, generating a high-concentration region relative to the sheet interior.

Based on this framework, we hypothesized that applying Wnt11-containing conditioned medium (CM) to migrating MDCK cells would generate a high-Wnt region near the leading edge and a relatively uniform low-Wnt region in the interior. To test this, we applied Wnt11 CM or parental CM (pCM) 8 h after confinement removal, when collective migration was established, and analyzed RP3 localization after 48 h. In cells treated with Wnt11 CM, RP3 exhibited a pronounced bias toward the side opposite the leading edge in a region slightly interior to the edge (100-500 μm) (Fig. 1A, B), whereas no such bias was observed with pCM (Fig. 1B). Quantification of polarity magnitude and polarity angle of RP3 confirmed that Wnt11 CM induces significant polarization compared to pCM (Fig. 1C, D). As an internal control, E-cadherin showed a uniform distribution even under conditions with Wnt11 CM (Fig. 1B, E), whereas RP3 polarity angles were significantly biased and differed from those of E-cadherin (Fig. 1F). This effect was recapitulated by addition of recombinant Wnt11 to the basal culture medium, indicating that Wnt11 has polarizing activity (fig. S2A).

**Fig. 1.**
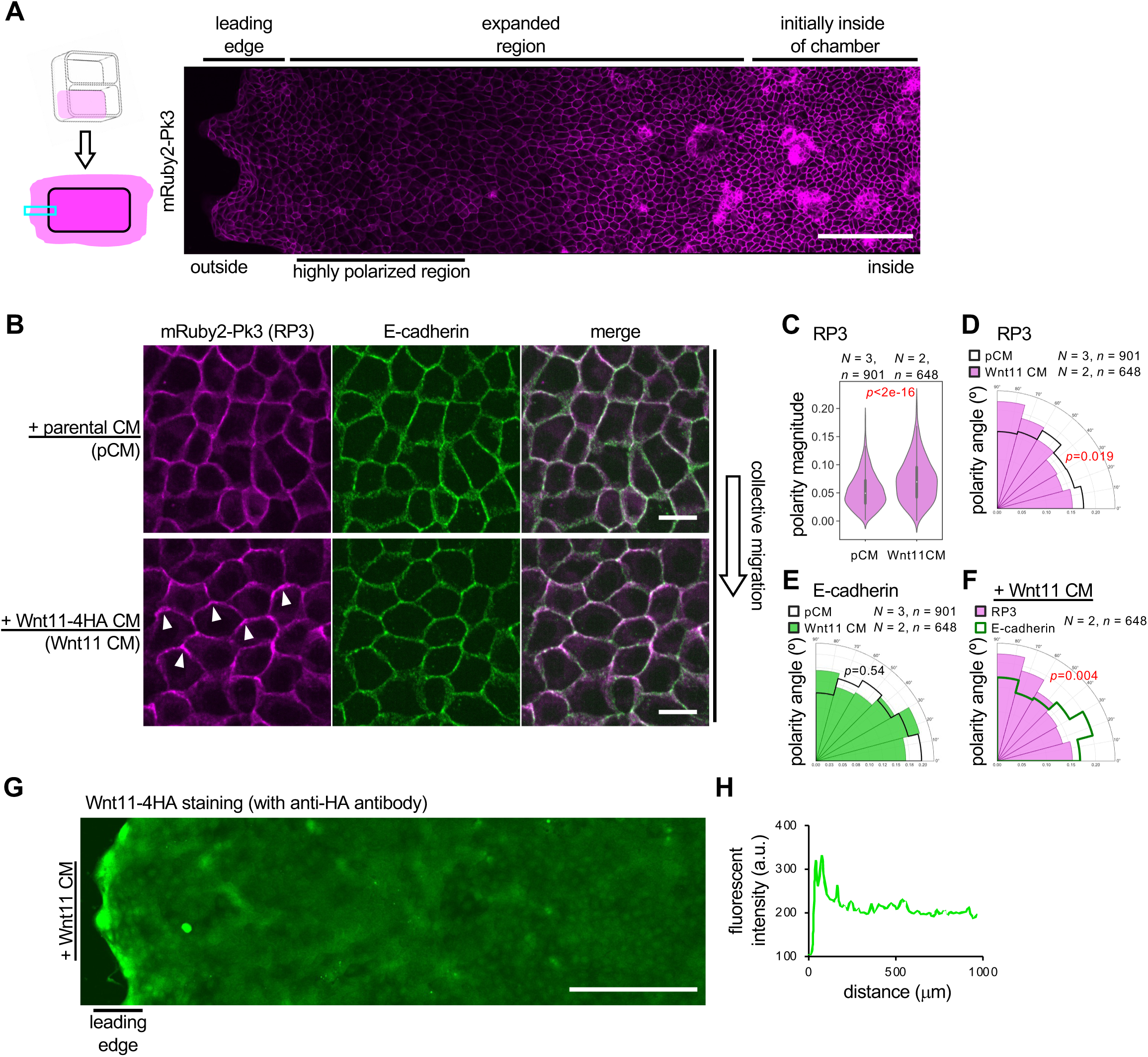
Wnt-dependent formation of planar cell polarity *in vitro*. **A**, *In vitro* polarization of mRuby2-Pk3 (RP3) by a combination of collective cell migration and Wnt11-4HA containing conditioned medium (Wnt11 CM). The observed area (cyan box) is indicated in the schematic at left; the region outlined in black denotes the seeding area inside the chamber, and cells collectively migrate out of this area after chamber removal. **B**, Wnt11 CM-dependent polarization of RP3. Regions slightly interior to the edge (100-500 μm) are shown. Arrowheads indicate planar polarized accumulation of RP3. **C-F**, Quantification of RP3 polarity with or without Wnt11 CM. Regions slightly interior to the edge (100-500 μm) are quantified. C, Polarity magnitude. D,E, Polarity angle with parental CM (pCM) or Wnt11 CM. F, polarity angle with Wnt11 CM (comparison between RP3 and E-cadherin in the same cells). Kuiper two-sample test (a circular analog of the Kolmogorov–Smirnov test) test was used for statistical analysis. **G**, **H**, Spatial distribution of Wnt11-4HA in the polarization condition. H, Wnt11-4HA was stained with anti-HA antibody, showing accumulation at the leading edge. I, quantification of fluorescent intensity of Wnt11-4HA staining along the outer-to-inner axis. Scale bars, 200 μm (A, G), 20 μm (B). Numbers of pictures (N) and numbers of cells (n) are as indicated. Representative data from two independent experiments are presented.

To understand how Wnt11 polarize cells, we next examined the distribution of Wnt11 protein itself. Using a C-terminally HA-tagged Wnt11 (Wnt11-4HA) in Wnt11 CM, we visualized its localization by anti-HA staining. Cells treated with pCM showed only background staining (fig. S2B), whereas Wnt11 CM produced strong staining near the leading edge and weaker, relatively uniform distribution in the interior (Fig. 1G, H, and fig. S2B). Thus, the predicted high-low Wnt distribution was indeed established by a combination of collective migration and Wnt11 CM, which is sufficient to reconstitute tissue wide coordination of PK3 polarity.

### Core PCP interdependence is recapitulated *in vitro*

A hallmark of PCP *in vivo* is mutual dependence among core PCP components. Therefore, we asked whether such interdependence is preserved in our *in vitro* PCP system using loss-of-function approaches. Among core PCP components expressed in MDCK II cells, we focused on *Vangl* genes (*Vangl1* and *Vangl2*), which comprise a relatively small number of homologs compared to other core PCP components. Published RNA-seq data indicated that *Vangl1* is the predominantly expressed isoform (*42*), so we generated Vangl1 knockout (KO) cells in an RP3-overexpressing background (Fig. 2A). As expected, membrane-associated staining of endogenous Vangl1 was abolished in these cells (Fig. 2B).

**Fig. 2.**
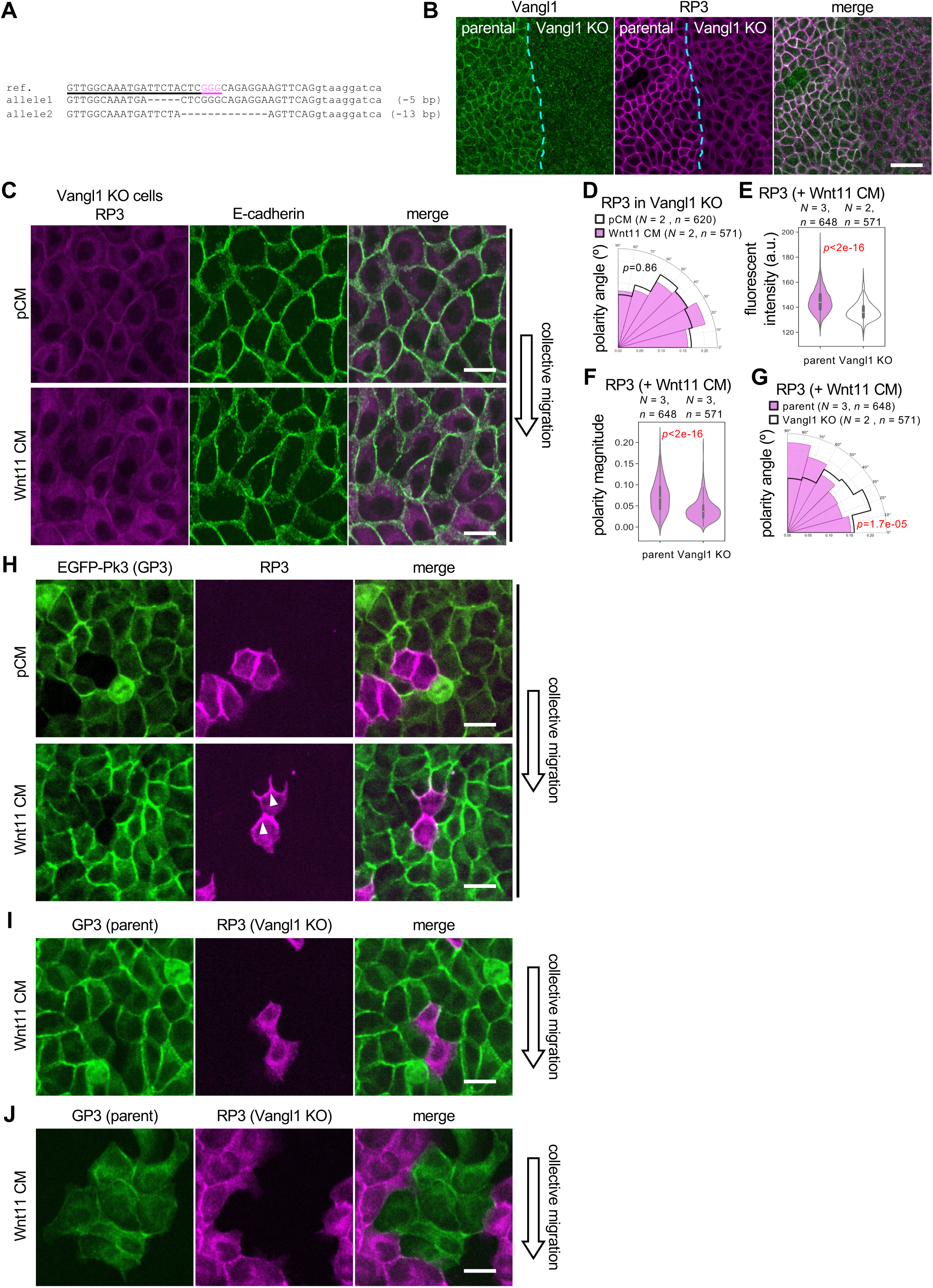
Cell-autonomous and non-cell-autonomous requirement of Vangl1 for vectorial polarization of Pk3. **A**, Genomic sequence of Vangl1 KO cells (clone 18D). Vangl1 KO cells were generated from RP3-overexpressing cells using CRISPR/Cas9 technique. The gRNA target sequence in exon 2 is underlined, and the PAM sequence is shown in magenta. Uppercase and lowercase letters indicate exon and intron sequences, respectively. Both alleles generated in-frame stop codons upstream of the coding sequence for the transmembrane domains. **B**, Loss of endogenous Vangl1 staining and reduced membrane localization of RP3 in Vangl1 KO cells. **C-G**, *In vitro* PCP formation was disrupted in Vangl1 KO cells. D, Conparison of polarity angle between pCM and Wnt11 CM. E, fluorescent intensity of membrane-associated RP3. F, Polarity magnitude of RP3 in parental cells and Vangl1 KO cells treated with Wnt11 CM. G, Conparison of polarity angle between parental Cells and Vangl1 KO cells treated with Wnt11 CM. **H**, Vectorial polarization of Pk3. Mixture of 95% EGFP-Pk3 (GP3)-overexpressing cells and 5% RP3-overexpressing cells were cultured in pCM or Wnt11 CM. Arrowheads indicate unipolar accumulations of RP3. See also fig. S3A for inverse ratio of GP3- and RP3-overexpressing cells. **I**, Cell-autonomous requirement of Vangl1. Mixture of 5% RP3-overexpressing Vangl1 KO cells and 95% GP3-overexpressing cells were cultured in Wnt11 CM. Unipolar accumulation of RP3 was not observed in Vangl1 KO cells. See also fig. S3B for the pCM condition. **J**, Non-cell-autonomous requirement of Vangl1. Mixture of 5% GP3-overexpressing cells and 95% RP3-overexpressing Vangl1 KO cells were cultured in the polarization condition. Unipolar accumulation of GP3 was not observed in parental cells. See also fig. S3C for the pCM condition. Scale bars, 40 μm (B), 20 μm (C, H, I). Numbers of pictures (N) and numbers of cells (n) are as indicated. Representative data from two independent experiments are presented.

In Vangl1 KO cells, RP3 membrane localization was reduced compared with parental cells under both maintenance and polarization conditions (Fig. 2B, E). Strikingly, RP3 failed to polarize in Vangl1 KO cells upon Wnt11 CM treatment (Fig. 2C, D). Compared with parental cells, RP3 polarity magnitude was significantly reduced in Vangl1 KO cells (Fig. 2F), and the distribution of polarity angles was also significantly different (Fig. 2G). These results demonstrate that interdependence among core PCP components is preserved in our *in vitro* system, as observed *in vivo* (*26, 43–46*).

Because conventional microscopy cannot resolve whether RP3 localizes to one or both sides of a cell junction, we mixed cells overexpressing EGFP-Pk3 (GP3) and RP3 at 95% and 5%, respectively, and performed the polarization assay with Wnt11 CM. Analysis of isolated RP3-expressing cells revealed that Pk3 localizes preferentially to the cell boundary opposite the leading edge (opposite the high-Wnt region) (Fig. 2H, see also fig. S3A for the reversed ratio of GP3- and RP3-expressing cells). These results demonstrate that these conditions can polarize Pk3 *in vitro* in a unipolar manner, as observed *in Xenopus* embryos (*26, 38*).

We next examined the cell autonomy of this effect. When a small fraction of RP3-overexpressing Vangl1 KO cells was mixed with a majority of GP3-overexpressing parental cells, RP3 was not polarized in Vangl1 KO cells even though neighboring parental cells exhibited polarization of GP3 (Fig. 2I and fig. S3B). Conversely, when a small number of GP3-overexpressing parental cells was mixed with a majority of RP3-overexpressing Vangl1 KO cells, GP3 polarization was also suppressed in parental cells (Figures 2J, S3A and S3C). Together, these results indicate that Vangl1 is required both cell-autonomously and non-cell-autonomously for asymmetric localization of Pk3 in this *in vitro* PCP system.

### Pk3 drives PCP reconstitution by modulating Vangl phosphorylation

In our *in vitro* PCP system, RP3 exhibits unipolar localization; thus, it serves as a robust readout of PCP. However, it remained unclear whether overexpressed RP3 acts merely as a reporter of intrinsic planar polarity, if any, of migrating MDCK cells as observed with polarized localization of Rac1, ppMLC, and the Golgi apparatus (*39, 40*) or whether it contributes functionally to PCP establishment. To address this question, we compared parental MDCK cells and RP3-overexpressing cells under polarization conditions (collective migration with Wnt11 CM). Immunostaining of endogenous Vangl1 revealed enhanced membrane localization in RP3-overexpressing cells compared to parental cells (Fig. 3A, B). Notably, Vangl1 exhibited significantly higher polarity magnitude in RP3-overexpressing cells than in parental cells upon Wnt11 CM treatment (Fig. 3C), but polarity angles did not differ significantly between them (fig. S4A). In addition, Vangl1 in Wnt11 CM-treated parental cells shows a slight, but significantly higher magnitude polarity than that in pCM-treated parental cells (Fig. 3D), while signaficant difference of the polarity angles of Vangl1 between pCM- and Wnt11 CM-treated cells was not observed (fig. S4B). These results indicate that RP3 is not simply a reporter, but is an essential component of PCP reconstitution, or at least, it enhances polarity of endogenous Vangl1.

**Fig. 3.**
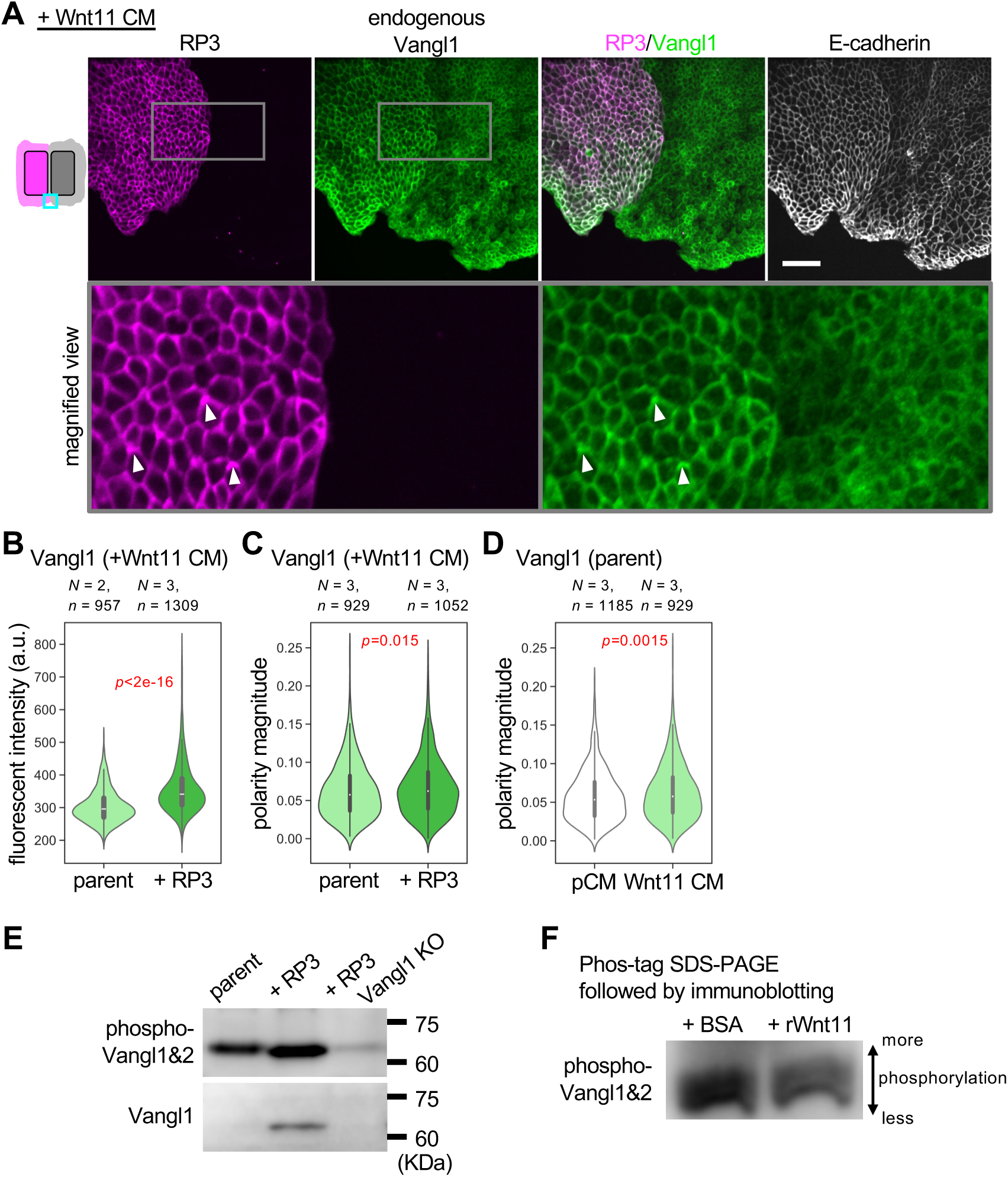
Overexpressed Pk3 facilitates polarization of endogenous Vangl1 via modulation of phosphorylation states of Vangl1. **A,** RP3 overexpression enhances plasma membrane localization of endogenous Vangl1. RP3-overexpressing cells and parental MDCK cells were seeded in the left and right regions, respectively, and cultured under polarization conditions with Wnt11 CM. **B**, Quantification of membrane-associated endogenous Vangl1 under polarization conditions. **C**, Polarity magnitude of endogenous Vangl1 in RP3-overexpressing cells and parental cells under polarization conditions. **D,** Polarity magnitude of endogenous Vangl1 in parental cells treated with pCM or Wnt11 CM. **E**, Immunoblotting. Lysate of parental, Pk3-overexpressing and Pk3-overexpressing Vangl1 KO cells at a fully confluent state, cultured in DMEM/10% FBS for 1 week were analyzed. **F**, Phos-tag SDS-PAGE (*48*) followed by immunoblotting was used to achieve phosphorylation-dependent segregation of Vangl1 (lower mobility corresponds to higher degree of phosphorylation). Bovine serum albumin (BSA) was used the vehicle control for recombinant Wnt11 (rWnt11, 2.0 μg/ml). See also fig. S5 for uncropped images of immunoblotting and images of PDVF membranes. Scale bars, 100 μm. Numbers of pictures (N) and numbers of cells (n) are as indicated. Representative data from two independent experiments are presented.

To investigate the mechanism that promotes polarization of Vangl1 in a Pk3-dependent manner, we examined the phosphorylation state of Vangl proteins. Phosphorylation of Vangl has been proposed as a central regulatory node in Wnt/PCP signaling (*23, 29*), directly modulating its interaction with Prickle (*26, 30*). Moreover, overexpressed Pk3 inhibits Vangl phosphorylation (*30, 47*). For this reason, we performed immunoblot analyses comparing parental cells and RP3-overexpressing cells. Phospho-Vangl1/2 (T78/S79/S82) bands were detected in both parental and RP3-overexpressing cells, whereas only weak bands were observed in RP3-overexpressing Vangl1 KO cells, likely reflecting trace amounts of Vangl2 expression (Fig.3E). In contrast, immunoblotting with an anti-Vangl1 revealed no detectable signal in parental cells, but a clear band in RP3-overexpressing cells, which was abolished in RP3-overexpressing Vangl1 KO cells, showing the specificity to Vangl1 (Fig.3E). This anti-Vangl1 antibody was raised against a recombinant N-terminal fragment of Vangl1 expressed in *E. coli*, which largely overlaps with known phosphorylation sites. We therefore speculate that this antibody preferentially recognizes non-phosphorylated Vangl1, as it failed to detect Vangl1 in parental cells by immunoblotting. These results suggest that Vangl1 is predominantly phosphorylated in parental cells, whereas RP3 overexpression induces a non-phosphorylated pool of Vangl1, consistent with the inhibitory effect of Pk3 on Vangl phosphorylation (*30, 47*). Thus, RP3 overexpression generates a cellular context in which phosphorylated and non-phosphorylated Vangl proteins coexist. This condition closely mirrors our findings in *Xenopus* embryos, where the coexistence of distinct phosphorylation states of Vangl is essential to drive asymmetric Pk3 localization (*26*).

In *Xenopus*, Wnt11 can induce phosphorylation of Vangl2 (*26*). Therefore, we also examined phosphorylation of Vangl proteins upon Wnt11 stimulation. Phos-tag SDS-PAGE (*48*) followed by immunoblotting revealed upper shifts of bands of phospho-Vangl proteins by addition of recombinant Wnt11 (Fig. 3F), suggesting increase of phosphorylation by Wnt11. Together, these results suggest that modulation of Vangl phosphorylation by Pk3 and Wnt11 is a key mechanism underlying PCP reconstitution *in vitro*.

### Dynamics of ERK signaling is involved in PCP formation

In collectively migrating MDCK cells, extracellular signal-regulated kinase (ERK) activity propagates as traveling waves from the leading edge and regulates migration dynamics (*40, 49*). In addition, FGF signaling, an upstream regulator of ERK signaling, has been implicated in PCP establishment in vertebrates (*24, 25*). These observations raised the possibility that ERK signaling may also contribute to PCP formation in our *in vitro* system. To test this idea, we inhibited ERK signaling using the MEK inhibitor PD0325901. Compared with the negative control (+DMSO), treatment with PD0325901 significantly reduced membrane-associated RP3 and RP3 polarity magnitude, although polarity angle was not significantly altered (Fig. 4A-C and fig. S6). These results suggest that ERK signaling contributes to PCP formation *in vitro*, at least in part by promoting membrane localization and polarized accumulation of RP3.

**Fig. 4.**
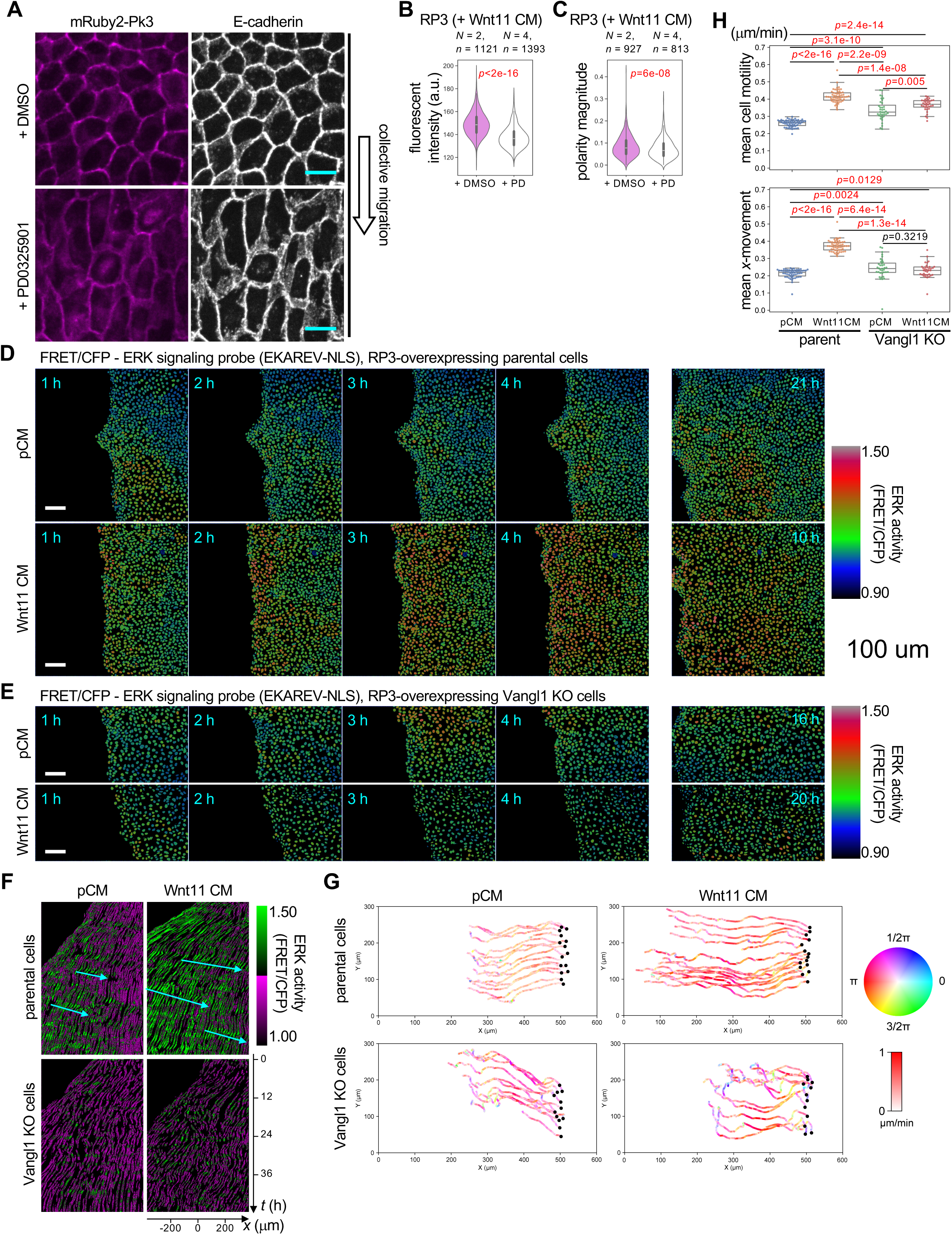
ERK signaling in *in vitro* PCP formation. **A-C**, Inhibition of ERK signaling by the MEK inhibitor PD0325901. DMSO (vehicle control) or PD0325901 (1 µM) was added under polarization conditions induced by collective migration and Wnt11 CM. **D, E**, ERK activity. FRET/CFP ratio images of the ERK activity probe EKAREV-NLS (*50*) in RP3-overexpressing parental cells (D) and Vangl1 KO cells (E) treated with pCM or Wnt11 CM. Images are shown at 1, 2, 3, and 4 h after the start of time-lapse imaging. The rightmost panels show the time point at which the leading edge reached the left edge of the imaging field. Images were acquired at 10-min intervals. **F**, *x*-*t* kymographs of ERK activity from the same data as in D and E. Cyan arrows indicate the rightward propagation of ERK activation waves. **G**, Cell trajectories. Cells located approximately 10 cell diameters from the leading edge at 1 h after the start of time-lapse imaging were tracked for 18 h. **H**, Mean cell motility and mean *x*-directional movement. Cells located approximately 10-15 cell diameters from the leading edge at 1 h after the start of time-lapse imaging were tracked for 18 h. Mean cell motility was defined as trajectory length divided by tracking duration, and mean *x*-directional movement (mean *x*-movement) as *x*-directional displacement divided by tracking duration. Scale bars, 20 µm (A), 100 µm (D, E). Numbers of pictures (N) and numbers of cells (n) are as indicated. Representative data from two independent experiments are shown.

To better understand the relationship between ERK signaling and PCP formation, we analyzed ERK dynamics by time-lapse imaging using the ERK activity probe EKAREV-NLS (*50*) under conditions with or without Pk3 polarization. In parental RP3-overexpressing cells treated with pCM, ERK activity propagated inward from the leading edge, as reported previously (Fig. 4D and movies S1 and S2). This inward propagation was also evident in an *x*-*t* kymograph (Fig. 4F). However, high ERK activity was only transient within individual cells, and its spatial spread was relatively limited. By contrast, in cells treated with Wnt11 CM, high ERK activity was more sustained and distributed over a broader area (Fig. 4D, F). In addition, the leading edge advanced at nearly twice the speed observed in pCM-treated cells (Fig. 4D). Consistent with this, cells in the region where polarity was observed moved faster under Wnt11 CM than under pCM (Fig. 4G). These distinct patterns of cell movement were consistent with the observed ERK activity dynamics.

We next examined ERK signaling and cell movement dynamics in Vangl1 KO cells (Fig. 4E-G). Strikingly, under both pCM and Wnt11 CM conditions, ERK activity waves propagating opposite to the direction of collective migration were abolished (Fig. 4F and movies S3 and S4). Cell trajectories also became markedly less linear, especially with Wnt11 CM (Fig. 4G). These findings suggest that Vangl1 is required for proper ERK wave propagation and for the coordinated collective migration. Quantification of cell movements further revealed that Wnt11 CM increased mean cell motility (defined as trajectory length divided by tracking duration) in both parental and Vangl1 KO cells. However, Wnt11 CM increased mean *x*-directional movement (*x*-movement; defined as *x*-directional displacement divided by tracking duration), which reflects collective migration, only in parental cells (Fig. 4H). These results suggest that Vangl1 is required to coordinate individual cell movements into directional collective migration.

Taken together, these findings suggest that Wnt11 alters ERK activity and cell motility dynamics, and that Vangl1 is required for their proper integration during directional collective migration associated with PCP formation. More broadly, these results point to a bidirectional relationship between Wnt/PCP signaling and ERK dynamics.

### Wnt11 provides directional cues, while tissue tension enables PCP establishment

In our *in vitro* PCP system, the combination of Wnt11 signaling and collective migration biases the unipolar component RP3 to the side opposite the leading edge. Because Wnt11 accumulation is highest at the leading edge, this orientation corresponds to distal side from regions of high Wnt11, consistent with the behavior of Pk3 observed in *Xenopus* embryos. However, it remained unclear whether directional cues are primarily dictated by Wnt11 signaling or by collective migration.

To distinguish these effects, we generated local sources of Wnt11 in the epithelial sheet by seeding islands of Wnt11-overexpressing cells prior to RP3-overexpressing cells and induced collective migration of RP3-expressing cells. Under these conditions, RP3 localization remained globally biased toward the side of each cell opposite the leading edge (Fig. 5A). However, at cell boundaries adjacent to Wnt11-overexpressing cells, membrane localization of Pk3 was locally reduced in a contact-dependent manner, closely resembling observations in *Xenopus* embryos (*26*) (Fig. 5A). Moreover, in cells located 2-3 cell diameters away from Wnt11-expressing cells, RP3 polarity was reoriented away from these local Wnt11 sources, recapitulating effects of Wnt5a overexpression on endogenous Vangl2 polarity in the *Xenopus* neural plate (*38*). These data suggest that Wnt11 provides local directional cues that can override global polarity.

**Fig. 5.**
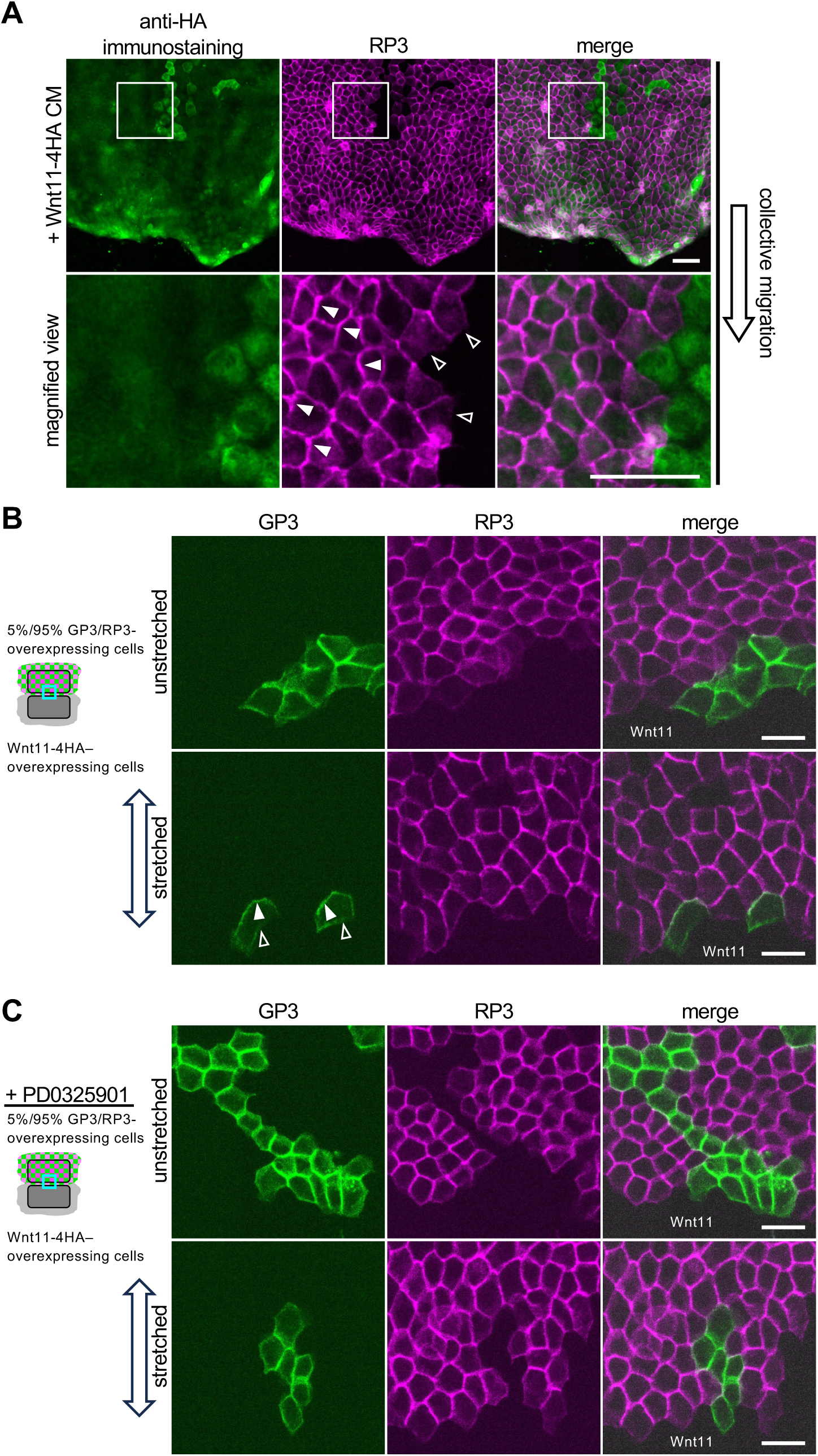
Tissue tension contributes to establishing polarity axis but not direction of Pk3 and local Wnt11 source biases the direction. **A**, Direction of RP3 polarity can be biased by local Wnt11 source. Wnt11-4HA–overexpressing cells (stained green) were seeded prior to RP3-overexpressing cells to make local Wnt11 sources. Wnt11 CM was also used as a permissive factor for RP3 polarization. Closed arrowheads indicate RP3 accumulation; open arrowheads indicate reduced RP3 localization. **B**, A local Wnt11 source together with tissue tension is sufficient to induce unipolar localization of Pk3. Cells were seeded as illustrated, and the imaged regions correspond to the boundaries between GP3- or RP3-overexpressing cells and Wnt11-4HA–overexpressing cells. Closed arrowheads indicate GP3 accumulation; open arrowheads indicate reduced GP3 localization. Wnt11-4HA–overexpressing cells correspond to regions lacking green and magenta fluorescence and are labeled “Wnt11” in the merged panels. **C**, MEK inhibition disrupts Pk3 polarization induced by a local Wnt11 source and tissue tension. Scale bars, 50 μm. Representative data from two independent experiments are presented.

We next asked whether collective migration per se is required, or whether the associated tissue tension is sufficient to establish polarity. To address this, we applied mechanical stretch to epithelial sheets in which Wnt11-overexpressing cells and Pk3-overexpressing cells were seeded adjacently. Under these conditions, cells did not undergo collective migration; nevertheless, upon application of tension, Pk3 was polarized away from Wnt11-overexpressing cells (Fig. 5B). This result indicates that tissue tension is sufficient to support polarity establishment, whereas collective migration itself is not strictly required. At least in this streching experiment, tension is inherently bidirectional and cannot define unipolarity by itself. Thus, these findings further suggest that directional information is specified by Wnt11, while tension contributes to establishing the polarity axis.

How do physical forces influence this biochemical polarity system? Mechanical stimuli activate ERK signaling (*40, 51*). Importantly, ERK inhibition also suppressed Pk3 polarization in the stretched condition, where collective migration was absent, but tissue tension was applied (Fig. 5C). Together, these results demonstrate that Wnt11 signaling and tissue tension act cooperatively to establish PCP *in vitro*, recapitulating key features observed in *Xenopus* embryos (*34*). Furthermore, ERK signaling is a potential mediator linking mechanical inputs to PCP establishment.

## Discussion

Here we establish a minimal epithelial system in which Wnt signaling and tissue tension together are sufficient to reconstitute key features of vertebrate planar cell polarity. In this system, tissue tension contributes to establishing an axis, but not its direction, while Wnt11 biases the unipolar localization of Pk3, together establishing tissue-wide vectorial polarity (Fig. 5). These findings move PCP reconstitution beyond tissue-scale alignment alone and identify a mechanistically tractable framework in which biochemical signaling and mechanical state can be experimentally separated and recombined.

Our results recapitulate key hallmarks of vertebrate PCP *in vivo*, including tissue wide vectorial polarity (Figs. 1 and 2), interdependence among core PCP components (Figs. 2 and 3), the requirement for Wnt signaling (Figs. 1 and 3), and regulation involving Vangl phosphorylation (Fig. 3). Notably, a high-low spatial pattern of Wnt11 combined with collective migration or tissue tension is sufficient to coordinate PCP over a long range (Figs. 1G and 5B). This provides a synthetic proof of concept for a model derived from our observations in *Xenopus* embryos: Wnt11 aligns PCP not by establishing a long-range gradient, but through local and reciprocal interactions with core PCP components (*26*).

ERK traveling waves have been intensively studied in collective migration (*40, 49*). In our system, ERK waves were more prominent under Wnt11 CM conditions, which accelerated collective migration, than under pCM conditions (Fig. 4D, F, H). Consistently, noncanonical Wnt signaling has previously been implicated in epithelial migration and wound repair (*52–54*). By contrast, the contribution of core PCP components to collective migration in epithelial cells has remained unclear, although they have been implicated in contact inhibition of locomotion in neural crest cells (*55*). We found that ERK traveling waves were abolished in Vangl1 KO cells (Fig. 4F). Consistent with this, Wnt11 CM no longer enhanced collective migration of Vangl1 KO cells, even though Wnt11 CM increased mean cell motility irrespective of Vangl1 (Fig. 4D, E, H). These results suggest that PCP contributes to the directional coordination of individual cell movements during collective migration, possibly by modulating ERK signaling dynamics. Furthermore, disruption of PCP by inhibition of ERK signaling (Fig. 4A-C) suggests a bidirectional regulatory relationship between PCP and ERK signaling.

The present system also clarifies the role of mechanics and the crosstalk between tensile force and biochemical signaling in PCP establishment. Although collective migration provided a robust and tractable context for polarity formation, migration itself was not strictly required once tissue tension was supplied externally (Fig. 5B). Mechanical stretch was sufficient to support Wnt11-dependent Pk3 polarization even in the absence of collective migration, indicating that tissue tension acts as a permissive or amplifying input rather than merely as a by-product of polarized movement. This finding recapitulates the cooperative action of Wnt signaling and tissue tension observed in the *Xenopus* neural plate (*34*). Inhibition of polarity by ERK signaling blockade under both migration-dependent and stretch-dependent conditions further suggests that ERK signaling may couple mechanical inputs to PCP establishment. In MDCK cells, collective migration and tissue tension activate ERK signaling (*40*). In some *in vivo* systems, such as mouse limb buds and *Xenopus* embryos, FGF signaling may also converge on ERK signaling (*24, 25*). Together, these observations support a two-input model in which local Wnt signals specify polarity direction, whereas tissue tension enables or stabilizes the polarized state with axial information, possibly via ERK signaling. Thus, they can substitute a body axis, which always underlies *in vivo* PCP.

While this work was in preparation, Wallach et al. reported synthetic reconstitution of planar polarity initiation in an engineered epithelial system and proposed collective migration as a symmetry-breaking cue (*56*). In their study, migration induced front-back enrichment of Celsr1 junctions and aligned tissue-scale axial polarity that tracked migration direction and speed, even in the absence of Vangl-dependent vectorial segregation. Our findings complement, but differ from this framework in several important respects. First, our readout is the unipolar localization of Pk3, a core PCP component, asymmetry of which depends on Vangl1, both cell-autonomously and non-cell-autonomously. Second, our system requires Wnt signaling, whereas tissue tension can substitute for migration itself. Third, our biochemical analyses link reconstituted polarity to the coexistence of distinct Vangl phosphorylation states generated by Pk3 overexpression. Therefore, we interpret the present system not as a migration-associated polarity state alone, but as a reductionist reconstruction of vertebrate PCP logic in which directional ligand input, mechanical competence, and interdependence of core PCP components are integrated.

Several limitations remain. Although our reconstitution recapitulates unipolar Pk3 asymmetry in a Wnt-and tension-dependent manner, as observed in the *Xenopus* neural plate, as well as the interdependence between Pk and Vangl (*34, 38*), it does not recapitulate convergent extension movements, which shape early vertebrate embryos (*5, 8*). Also, endogenous Celsr- and Frizzled-based intercellular complexes have not yet examined. Nevertheless, these findings illustrate how a developmental control system that normally operates *in vivo* can be reduced to experimentally controllable biochemical and mechanical inputs, providing a synthetic framework for dissecting PCP, drug screening, and understanding Wnt/PCP signaling.

## Materials and Methods

### Cell culture

MDCK II cells were provided by Dr. Masayuki Murata (The University of Tokyo) and cultured in low-glucose DMEM (Nissui/Shimadzu, #05919) supplemented with 10% fetal bovine serum (FBS) (*57*) in a humidified incubator at 37°C with 5% CO₂.

### Preparation of conditioned media and *in vitro* PCP formation assay

Parental or Wnt11 conditioned medium (CM) was prepared as follows. Parental or Wnt11-4HA–overexpressing MDCK cells were cultured in 10-cm dishes. After the cells reached full confluence, the medium was replaced with fresh DMEM supplemented with 10% FBS, and CM was collected 60 h later. Cell debris was removed by sequential centrifugation at 1,000 rpm for 5 min and 3,000 rpm for 10 min. The final supernatant was collected, aliquoted, snap-frozen in liquid nitrogen, and stored at −80°C. Immediately before use, frozen CM was thawed in a 37°C water bath with continuous gentle mixing.

For the *in vitro* PCP formation assay, cells were seeded into each chamber of a Culture-Insert placed in a 35-mm µ-Dish (Ibidi; 2-well, #81176, or 3-well, #80366). For each chamber, 70 µl of cell suspension in DMEM supplemented with 10% FBS at 8 × 10^5^ cells/ml was added and incubated for 16 h. The insert was then removed, and the medium was replaced with fresh DMEM containing 10% FBS. After 8 h, the medium was replaced with either parental or Wnt11 CM. After 24 h, the conditioned medium was replaced with fresh parental or Wnt11 CM, respectively, and cells were fixed 24 h later and processed for immunostaining. For experiments with recombinant Wnt11, recombinant human Wnt11 (R&D Systems, 6179-WN) was reconstituted at 80 μg/ml in 0.2% bovine serum albumin (BSA) in PBS and added to DMEM supplemented with 10% FBS at a final concentration of 2.0 μg/ml in place of Wnt11 CM.

### Molecular biology

The following genes were PCR-amplified and subcloned into the mammalian expression vector pCA Nw-Sal-Flag (*58*) to establish stable transfectants: EGFP-Xpk3 (*38*), mRuby2-Xpk3 (*26, 34*), Xwnt11-4HA (this study), secreted mVenus (*41*), and morphotrap (*59*). *Canis lupus familiaris* Vangl1 was PCR-amplified and subcloned into pCA H-Sal-Flag (*58*).

Stable transfectants were established by transfecting the plasmids using Lipofectamine LTX with PLUS reagent (Thermo Fisher Scientific, #A12621) according to the manufacturer’s instructions. Cells were selected with 500 µg/ml G418 (Nacalai Tesque, #09380-44), 100 µg/ml hygromycin (Nacalai Tesque, #09287-84), or 10 µg/ml blasticidin (InvivoGen, ant-bl-1). To establish stable EKAREV-NLS transfectants (*50*), pPBbsr2-EKAREV-NLS and pHyPBase (*60*) were co-transfected. Multiple surviving clones were isolated, and transgene expression was confirmed by immunostaining and confocal microscopy. Multiple clones were isolated for each construct, and all clones showed consistent phenotypes.

Vangl1 KO cells were generated by electroporating RP3-overexpressing cells with Cas9-gRNA RNP complexes as previously described (*61*). The target sequence was as follows, with the PAM sequence underlined: 5’-GTTGGCAAATGATTCTACTCGGG-3’. KO cells were screened by immunostaining in glass-bottom 96-well plates (Corning, #4580). Mutations in Vangl1 KO cells were confirmed by DNA sequencing of PCR fragments spanning the target sequence.

### Cell stretching experiments

Cells were seeded in a Culture-Insert 2 Well (Ibidi, #80209) placed on a stretching chamber (Strex, SC-0040 or SC-0044) precoated with bovine fibronectin (Sigma, F1141). Twenty-four hours after seeding, the Culture-Insert was removed, and the medium was replaced with fresh DMEM containing 10% FBS. Cells were then incubated for an additional 24 h. The medium was subsequently replaced with Wnt11 CM, and cells were incubated for 24 h before stretching. Chambers were stretched by 20% over 1 h using a manual stretcher (Strex, ST-0040 or ST-0100). After an additional 20 min, cells were fixed and analyzed. Where indicated, PD0325901 was added at a final concentration of 1 µM 1 h before stretching.

### Fluorescence image acquisition

Fluorescence images were acquired using either a spinning-disk confocal microscope (Andor Dragonfly 200 combined with an Olympus IX83; objective, UPLXAPO20X, 20×/0.80 NA) or a laser-scanning confocal microscope (Leica TCS SP8; objective, HC PL APO 40×/1.10 W CORR, water immersion), except for ERK activity analysis.

### ERK activity analysis

ERK activity analysis was performed using cells prepared in the same manner as for the *in vitro* PCP formation assay. Cells expressing EKAREV-NLS (*50*) were imaged with an IX83 inverted microscope (EVIDENT) equipped with an sCMOS camera (ORCA-Fusion BT; Hamamatsu Photonics), a spinning disk confocal unit (CSU-W1; Yokogawa Electric Corporation), and diode lasers. Images were acquired using an air/dry objective lens (UPLXAPO20X, 20×/0.80 NA). The excitation laser and fluorescence filter settings were as follows: excitation at 445 nm, dichroic mirror DM 445/514/640, and emission filters of 482/35 for CFP and 525/50 for FRET. The microscope system was controlled using MetaMorph software (version 7.10.3). During imaging, cells were maintained in a stage incubator (STXG-IX3WX; Tokai Hit) at 37°C under 5% CO2.

### Immunostaining

MDCK cells were fixed with MEMFA (0.1M MOPS, pH 7.4, 2 mM EGTA, 1mM MgSO_4_, 3.7% formaldehyde). After fixation, the cells were washed once with PBS and once with TBT (1x TBS, 0.2% BSA, 0.1% Triton X-100), and blocked with TBTS (TBT with 10% heat-immobilized FBS [70°C, 40 min]) (RT, 30 min, or 4°C 1-2 h). All cells were incubated with the primary antibodies diluted in Can Get Signal immunostain Immunoreaction Enhancer Solution A (NKB-501, TOYOBO), or B (NKB-601, TOYOBO), or TBTS (RT, 1 h), washed three times with TBT, and incubated with secondary antibodies diluted in TBTS (RT, 1 h). To reduce punctate staining artifacts, diluted antibodies were centrifuged at 4°C and 20,000 × g for 15 min, and the resulting supernatants were used for immunostaining. Finally, the cells were washed three times with TBT and mounted using ProLong Glass antifade mountant (Invitrogen, P36980). Primary antibodies and their dilutions were as follows: anti-Vangl1 (Sigma-Aldrich, HPA025235, 1/200), anti-E-cadherin (hybridoma culture supernatant of clone ECCD-2 (RIKEN BRC, RCB5259), prepared in-house, 1/20), anti-HA (MBL, #561, 1/200). Secondary antibodies and their dilutions were as follows: goat polyclonal anti-rabbit IgG Alexa Fluor 488 (Invitrogen, A11008, 1/500), goat polyclonal anti-rat IgG Alexa Fluor 647 (Invitrogen, A21247, 1/500).

### Image analyses

All measurements of signal intensity were performed using Fiji/ImageJ (v1.54f or v1.54p). For quantification of polarity angle and magnitude, cell segmentation masks were generated as follows. First, E-cadherin images were Gaussian-blurred (σ = 5). Background subtraction was then performed using the rolling-ball algorithm with a radius of 50 pixels. The images were binarized using a threshold range of 4–65,535 and then inverted. The resulting mask images were manually corrected using the Fiji/ImageJ plugin TissueAnalyzer (*62*). Polarity angles and magnitudes were quantified using QuantifyPolarity2.0 with the Fourier method (*63*). For measurement of signal intensities on the membrane in Figures. 2E and 3B, ROIs for individual cells were generated from E-cadherin images using Cellpose3 with the Cyto3 model (*64*). Intensity profiles were collected along 5-pixel-wide lines using an in-house Fiji/ImageJ macro.

### Immunoblotting

Immunoblotting was performed using a standard protocol as described previously (*65*). Briefly, samples were mixed with 2× SDS-PAGE sample buffer containing 5% (w/v) SDS, 17.6% (w/v) glycerol, 0.063 M Tris-HCl (pH 6.8), 10% (v/v) 2-mercaptoethanol, and 0.01% (w/v) bromophenol blue, heated at 95°C for 10 min, and loaded onto 10% polyacrylamide gels for electrophoresis. Proteins were then transferred to PVDF membranes, which were blocked with 5% skim milk before incubation with primary antibodies.

The following primary antibodies were used: anti-phosphorylated Vangl1/2-T78/S79/S82 (Abclonal, AP1206, rabbit monoclonal antibody, 1/2000) and anti-Vangl1 (Sigma-Aldrich, HPA025235, rabbit polyclonal antibody, 1/2000). For detection, HRP-conjugated anti-rabbit IgG (Jackson ImmunoResearch Laboratories, #111-035-144) and Chemi-Lumi One Super (Nacalai Tesque, #02230-14) were used.

For Phos-tag SDS-PAGE, 10% SuperSep Phos-tag precast gels containing 50 µmol/l Phos-tag (FUJIFILM Wako Pure Chemical) were used to separate phosphorylated proteins. Before transfer to PVDF membranes, the gels were washed three times in transfer buffer containing 10 mM EDTA, according to the manufacturer’s instructions.

### Statistical analyses

Sample sizes were determined empirically. The Kuiper two-sample test, a circular analog of the Kolmogorov–Smirnov test, was used to analyze polarity angle distributions. All other statistical analyses were performed using the Wilcoxon rank-sum test, with *p* values adjusted for multiple comparisons by Holm’s method.

## Acknowledgements

We thank Drs. Mototsugu Eiraku, Danelle Devenport, and Kazuhiro Aoki for discussion; Dr. Sergei Y. Sokol for cDNAs of EGFP-Pk3; Dr. Markus Affolter for morphotrap cDNA; Dr. Kosuke Yusa for pHyPBase; Drs. Toshihiko Fujimori and Mikio Furuse for technical support; Drs. Ritsuko Takada and Shinji Takada for discussion and technical support, and Dr. Steven D. Aird for editing the manuscript.

## Author contributions

Conceptualization: TO, YM; Methodology: SM, MS, SH, TO, YM

Software: MS;

Validation: SM, MS;

Formal analysis: SM, MS, SH, YM

Investigation: SM, MS, SH, YM

Resources: SM, TO, YM;

Data curation: SM, MS, SH, YM

Writing - original draft: YM;

Writing - review & editing: SM, MS, SH, TO

Visualization: SM, MS, SH;

Supervision: TO, YM

Project administration: YM;

Funding acquisition: TO, YM.

## Data and materials availability

All data needed to evaluate the conclusions in the paper are present in the paper and/or the Supplementary Materials.

## Supplementary Materials

figs. S1 to S6, movies S1-S4

## Supplementary Materials

### Supplementary Figures

**fig. S1.**
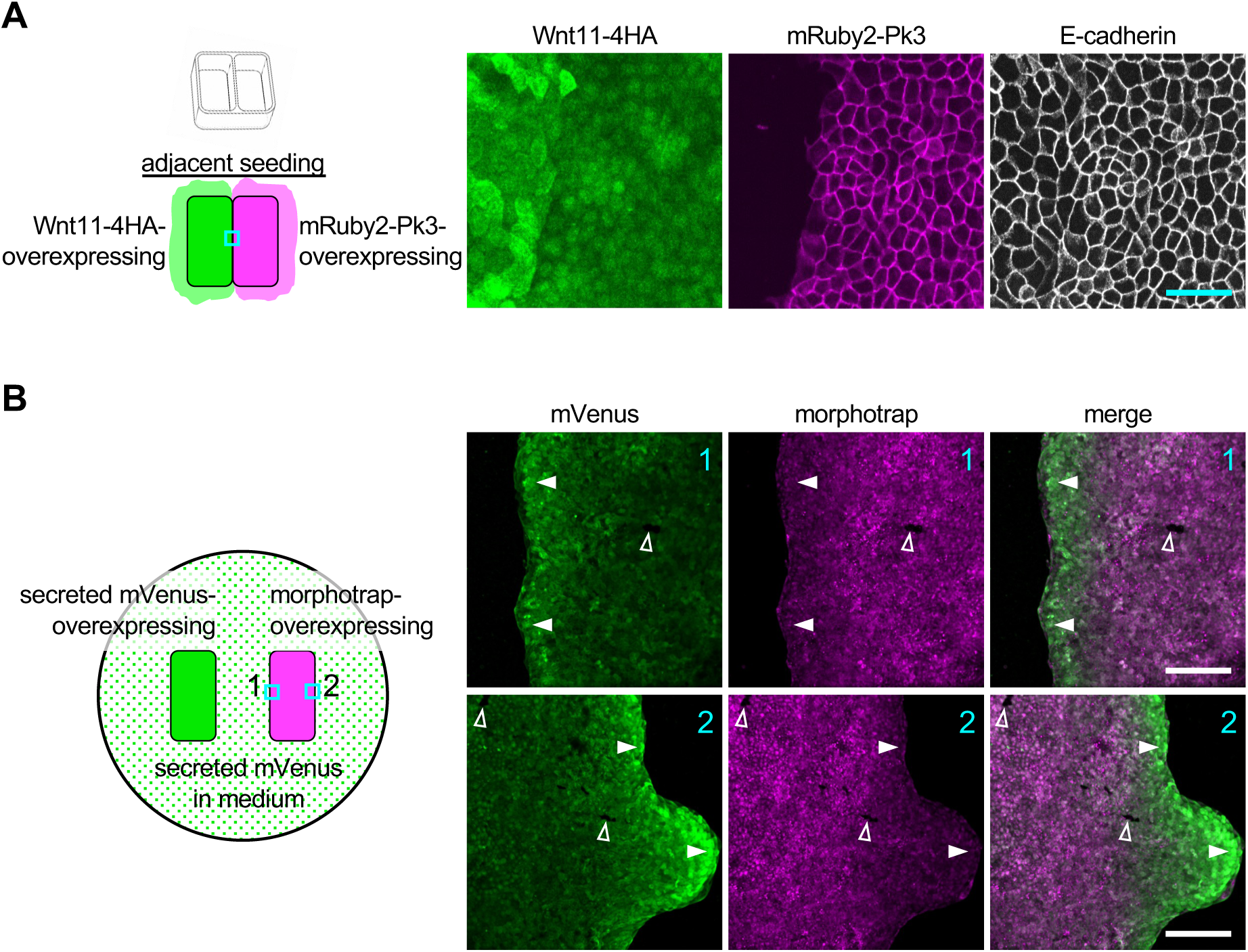
Adjacent positioning of Wnt11- and Pk3-overexpressing cells were not sufficient for planar polarization of Pk3. **A**, Wnt11-4HA– and mRuby2-Pk3–overexpressing cells were seeded adjacently using a two-well chamber. After chamber removal, the two cell populations came into contact along a straight boundary, which was typically established by 24 h. Twelve hours after chamber removal, the medium was replaced with Wnt11-4HA conditioned medium (CM), and the cells were cultured for an additional 48 h before fixation and immunostaining. The boundary region is shown. Unlike the polarized localization observed upon adjacent expression of Wnt11 and Pk3 in *Xenopus*, Pk3 polarization was not detected under these conditions. **B**, Accumulation of secreted mVenus at the leading edge of morphotrap-overexpressing cells. Morphotrap is a membrane-tethered anti-GFP nanobody (*59*). Secreted mVenus-overexpressing cells and morphotrap-overexpressing cells were seeded separately in the two outer wells of a three-well chamber, leaving the middle well empty. After the cells reached confluence, the chamber was removed, and live imaging was performed 48 h later. Secreted mVenus accumulated at both the left and right leading edges (closed arrowheads), suggesting that it diffused throughout the dish. Open arrowheads indicate cells that had lost morphotrap expression and no longer accumulated secreted mVenus, indicating that mVenus accumulation depends on morphotrap. Scale bars: 100 μm. Representative data from two independent experiments are presented.

**fig. S2.**
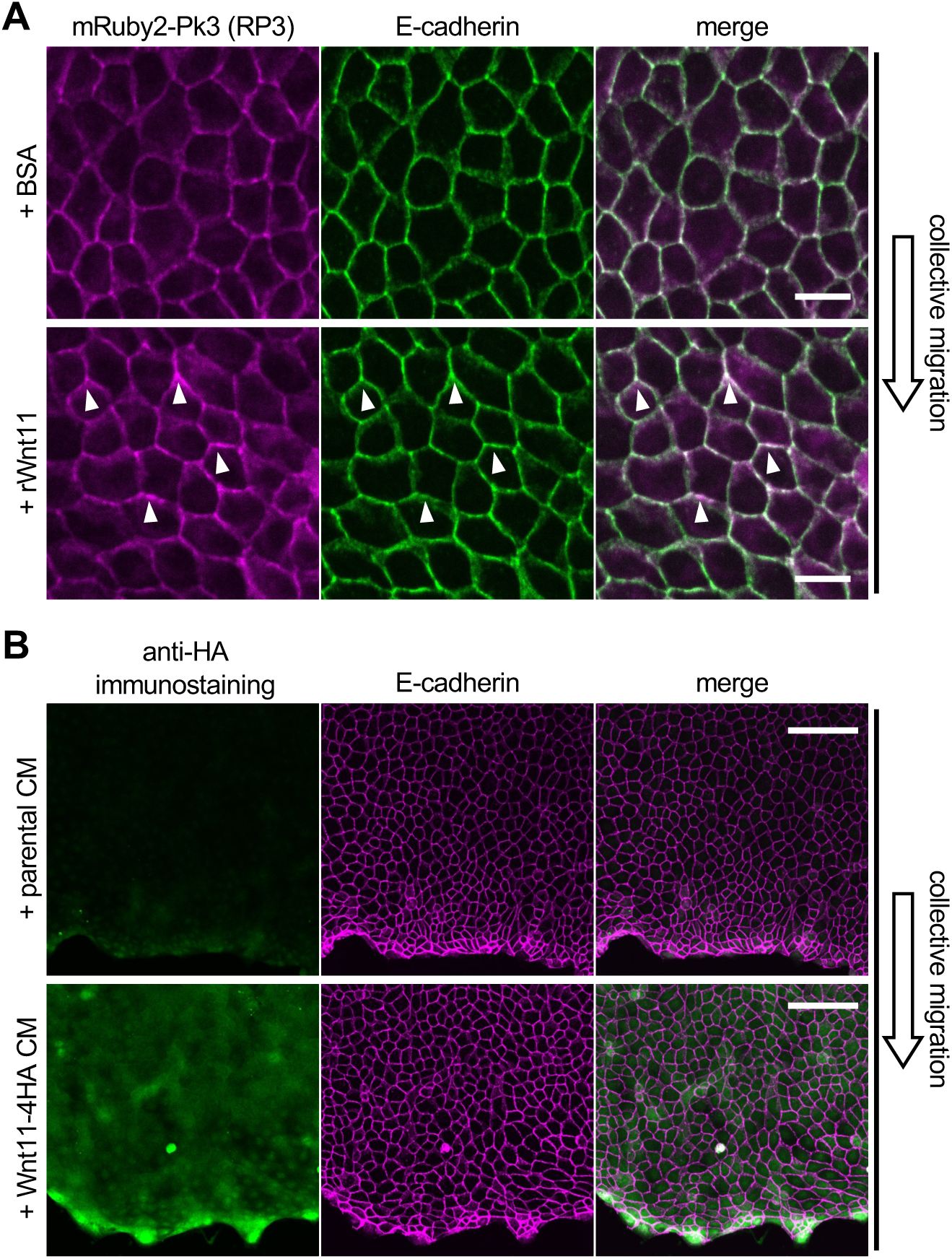
RP3 polarization by recombinant Wnt proteins and immunostaining of Wnt11-4HA. **A**, Addition of recombinant Wnt11 induced RP3 polarization in collectively migrating cells. Bovine serum albumin (BSA) was used the vehicle control for recombinant Wnt11 (rWnt11, 2.0 μg/ml). Arrowheads indicate polarized accumulation of RP3. **B**, Visualization of Wnt11-4HA protein. RP3-overexpressing cells were treated with parental CM or Wnt11-4HA CM for 48 h, during collective migration. Immunostaining with anti-HA antibody specifically detected Wnt11-4HA, with strong accumulation at the leading edge. Scale bars, 20 μm (A), 100 μm (B). Representative data from two independent experiments are presented.

**fig. S3.**
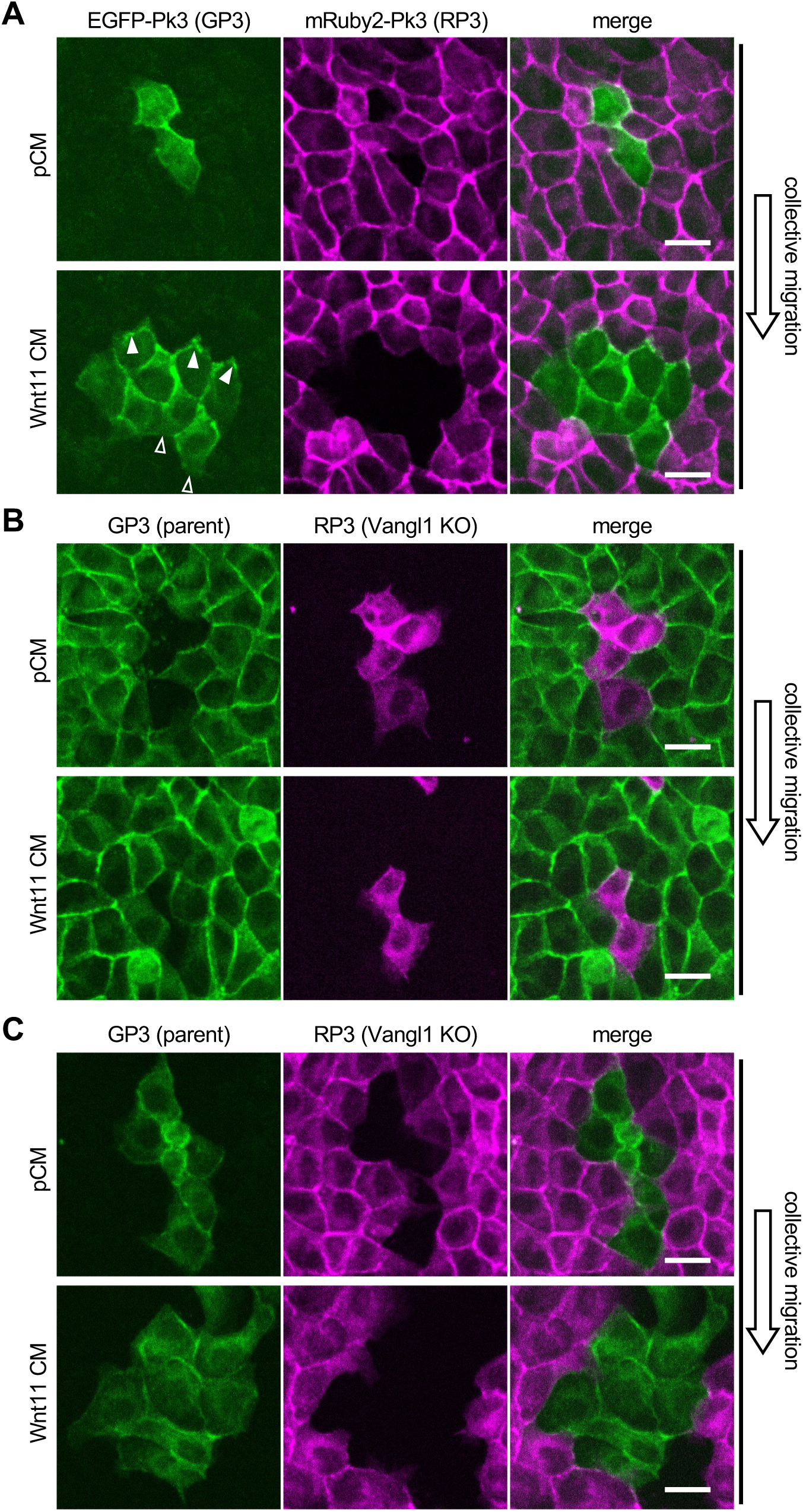
Mosaic analyses with Vangl1 KO cells. **A**, Mixture of 5% EGFP-Pk3 (GP3)-overexpressing cells and 95% RP3-overexpressing cells were cultured in pCM or Wnt11 CM. Closed and open arrowheads indicate accumulations and absence of GP3, respectively. **B**, Mixture of 95% GP3-overexpressing parental cells and 5% RP3-overexpressing Vangl1 KO cells were cultured in pCM or Wnt11 CM. Data labeled as “Wnt11 CM” are identical to those in Fig. 2I. **C**, Mixture of 5% GP3-overexpressing parental cells and 95% RP3-overexpressing Vangl1 KO cells were cultured in pCM or Wnt11 CM. Data labeled as “Wnt11 CM” are identical to those in Fig. 2J. Scale bars, 20 μm. Representative data from two independent experiments are presented.

**Fig. S4.**
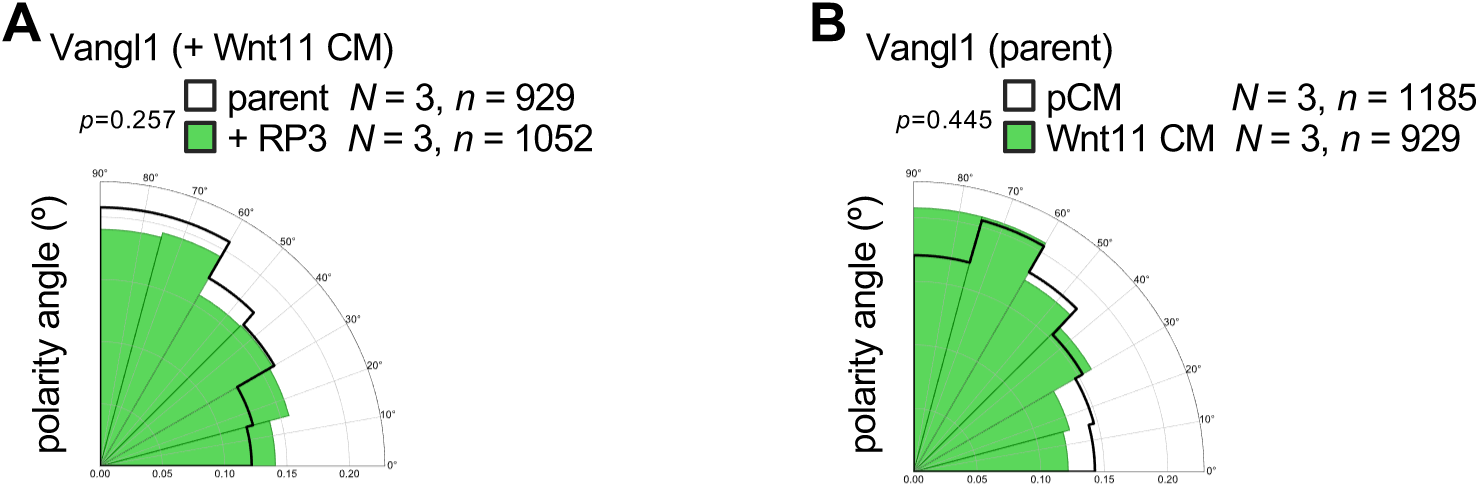
Polarity magnitude of endogenous Vangl1. **A**, Comparison between parental and RP3-overexpressing cells treated with Wnt11 CM. **B**, Comparison of parental cells treated with pCM or Wnt11 CM. Numbers of pictures (N) and numbers of cells (n) are as indicated.

**fig. S5.**
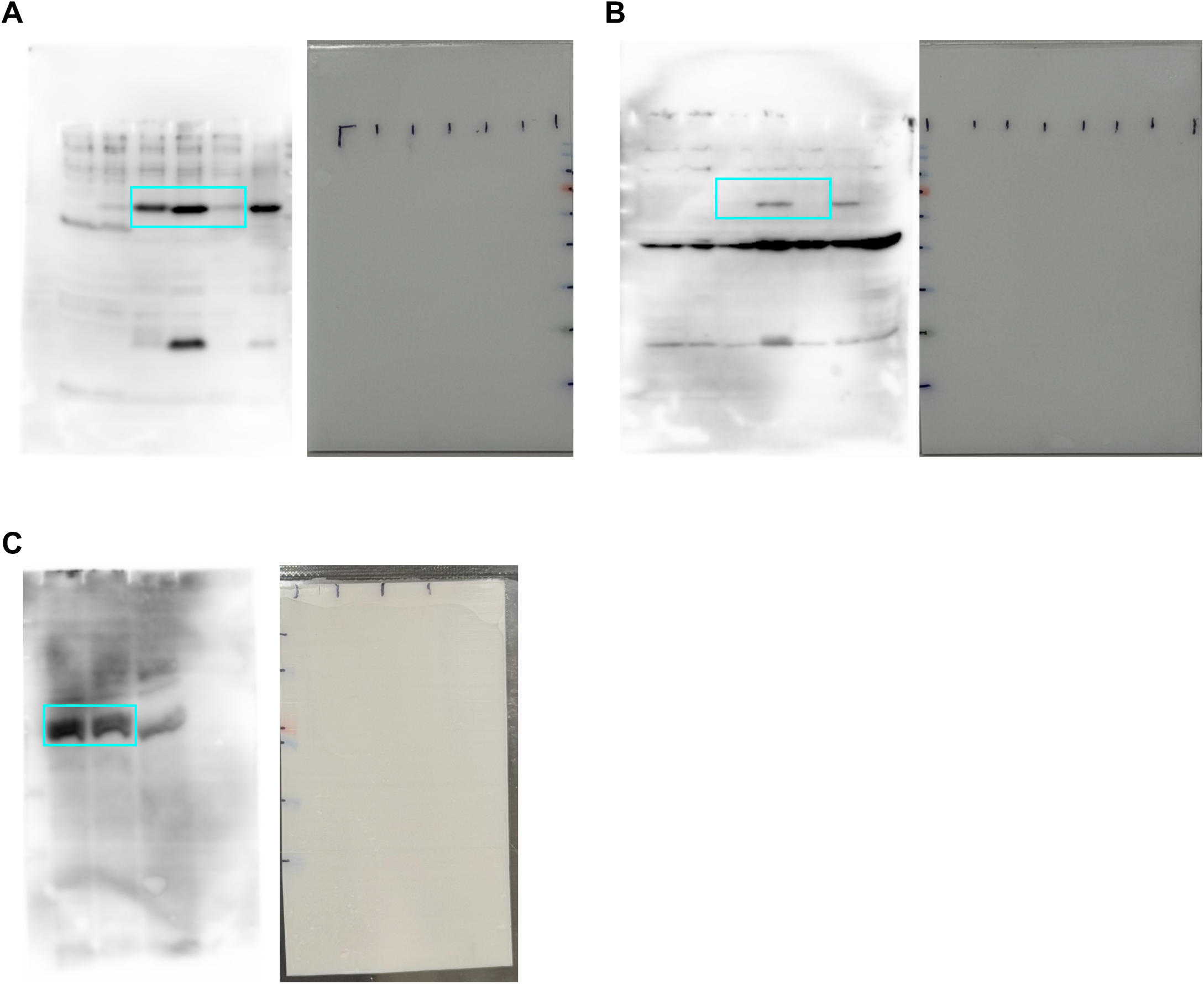
Uncropped images of immunoblotting and images of PDVF membranes. Regions shown in the main figures are boxed. A, for Fig 3E, “phospho-Vangl1&2”; B, for Fig 3E, “Vangl1”; C for Fig. 3F.

**fig. S6.**
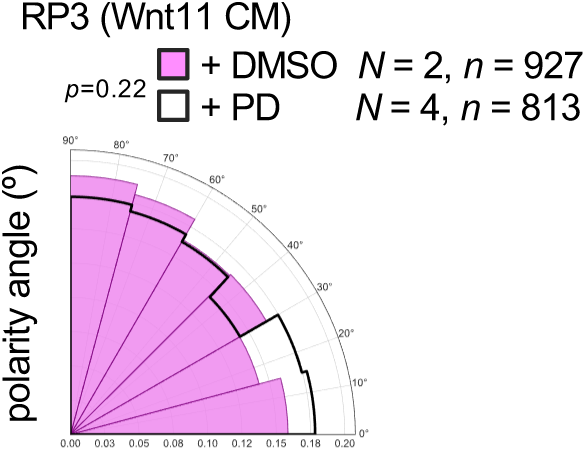
Polarity angle of RP3 in polarization conditions with or without PD0325901. DMSO (vehicle control) or PD0325901 (1 µM) was added under polarization conditions induced by collective migration and Wnt11 CM. Numbers of pictures (N) and numbers of cells (n) are as indicated.

### Supplementary Movies

**movies S1-S4. Dynamics of ERK activity in collectively migrating parental and Vangl1 KO cells treated with pCM or Wnt11 CM.**

FRET/CFP ratio images of RP3-overexpressing parental cells treated with pCM (movie S1) or Wnt11 CM (movie S2), and RP3-overexpressing Vangl1 KO cells treated with pCM (movie S3) or Wnt11 CM (movie S4). Images were acquired at 10-min intervals for 48 h. In the movies, 1 s corresponds to 2 h of real time. The same pseudocolor lookup table (LUT) used in Figures 4D and 4E was applied.

